# Discovery of a polybrominated aromatic secondary metabolite from a planctomycete points at an ambivalent interaction with its macroalgae host

**DOI:** 10.1101/723874

**Authors:** Fabian Panter, Ronald Garcia, Angela Thewes, Nestor Zaburannyi, Boyke Bunk, Jörg Overmann, Mary Victory Gutierrez, Daniel Krug, Rolf Müller

## Abstract

The roles of the majority of bacterial secondary metabolites, especially those from uncommon sources are yet elusive even though many of these compounds show striking biological activities. To further investigate the secondary metabolite repertoire of underexploited bacterial families, we chose to analyze a novel representative of the yet untapped bacterial phylum *Planctomycetes* for the production of secondary metabolites under laboratory culture conditions. Development of a planctomycetal high density cultivation technique in combination with high resolution mass spectrometric analysis revealed Planctomycetales strain 10988 to produce the plant toxin 3,5 dibromo p-anisic acid. This molecule represents the first secondary metabolite reported from any planctomycete. Genome mining revealed the biosynthetic origin of this doubly brominated secondary metabolite and a biosynthesis model for the com-pound was devised. Comparison of the biosynthetic route to biosynthetic gene clusters responsible for formation of polybrominated small aromatic compounds reveals evidence for an evolutionary link, while the compound’s herbicidal activity points towards an ambivalent role of the metabolite in the planctomycetal ecosystem.

## 1 Introduction

Bacterial secondary metabolism has long been a source of chemically diverse and biologically active natural products.^1,2^ In fact, large numbers of biologically active entities have been isolated from extensively screened phyla such as actinobacteria, firmicutes and proteobacteria.^3–5^ To establish alternative sources, natural products research is increasingly focusing on taxa that have been less exploited to date, but show potential for production of secondary metabolites according to the presence of secondary metabolite biosynthesis gene clusters (BGCs) in their genomes.^2^ This strategic shift towards new producers increases chances for the discovery of novel bioactive secondary metabolite scaffolds that are chemically distinct from the scaffolds found in previously screened bacteria. While it has long been stated that phylogenetically distant species have a more distinct secondary metabolism, recent comprehensive secondary metabolome studies were able to validate this claim.^6,7^ Accordingly, it is now widely recognized that there is an urgent need to scrutinize novel bacterial taxa, alongside the use of sensitive mass spectrometry and varied cultivation conditions to unearth novel natural products from bacterial secondary metabolomes.^8^ Planctomycetes represent an underexploited phylum of bacteria in terms of their secondary metabolite potential.^9^ Although planctomycetes have been already discovered in 1924, until now no secondary metabolite of planctomycetal origin has been reported.^10^ This bleak picture is in clear contrast to previous *in-silico* genome analysis that suggested planctomycetes to contain a significant number of secondary metabolite biosynthetic gene clusters (BGCs).^9,11^ In this work, we describe the first secondary metabolite from any planctomycete including its structural characterization and biosynthesis, whereas its biological activity sheds light on the putative ecological role within the planctomycete’s natural habitat.

## 2 Results and Discussion

### 2.1 Cultivation of *Planctomycetales* strain 10988

In order to investigate the biosynthetic capacity of uncommon and underexploited bacteria, we set out to isolate new strains from marine sediment samples. Our efforts revealed a swarming, rose colored bacterial isolate that was designated as strain 10988 (see Figure 1).

**Figure 1.**
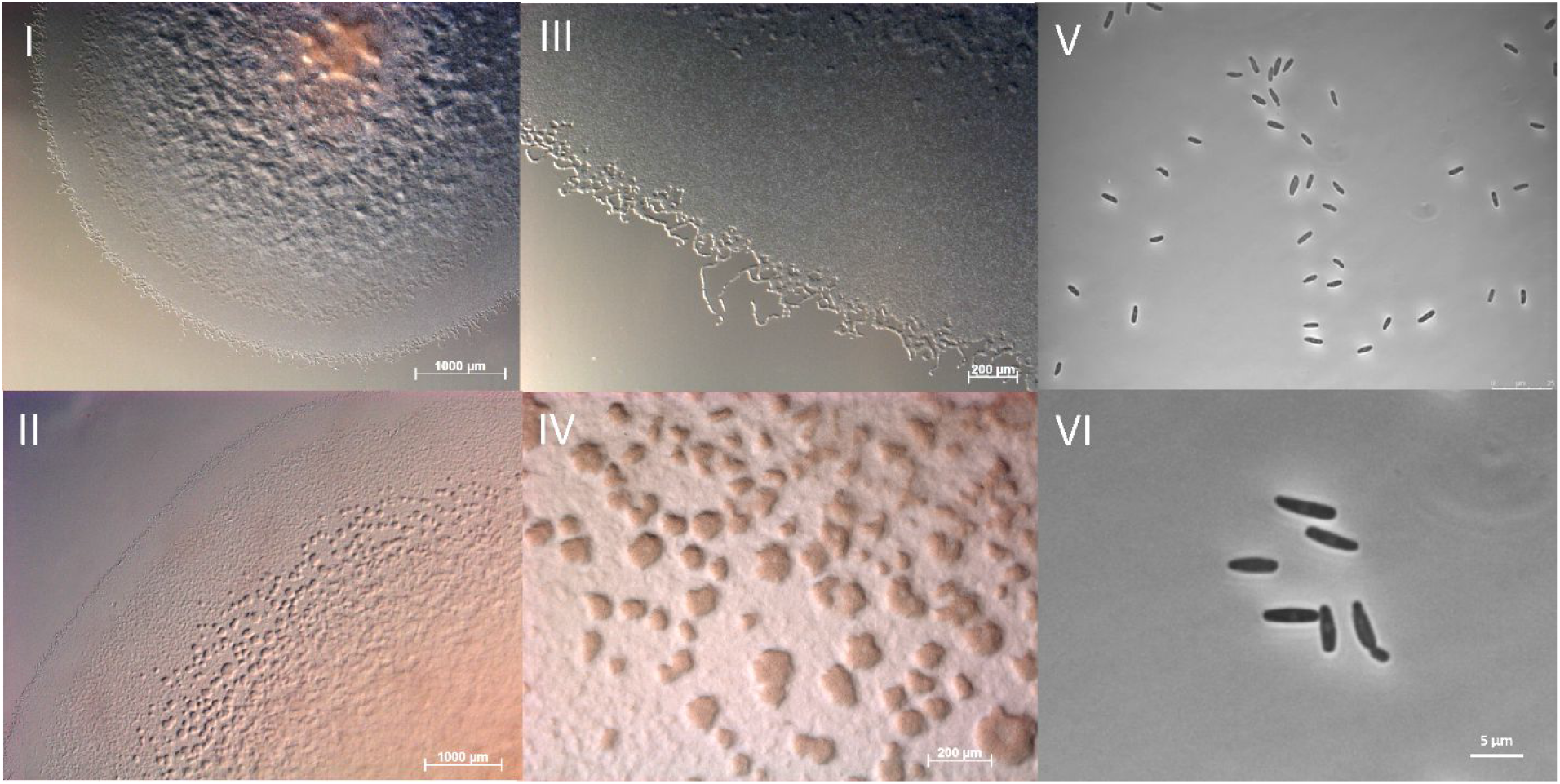
*Growth characteristics of* Planctomycetales *strain 10988 on solid medium displaying swarming (I and III), fruiting body like aggregate formation (II to IV) and phase contrast microscopy images of single cells from swarm (V) and from fruiting body like aggregates (VI).*

16S rRNA-gene based phylogenetic analyses revealed the bacterium to belong to the order *Planctomycetales*, while being genetically distant from previously characterized planctomycetal genera such as *Thermogutta, Thermostilla, Pirellula, Rhodopirellula* and *Blastopirellula* (see Figure 2).^9,12–14^

**Figure 2.**
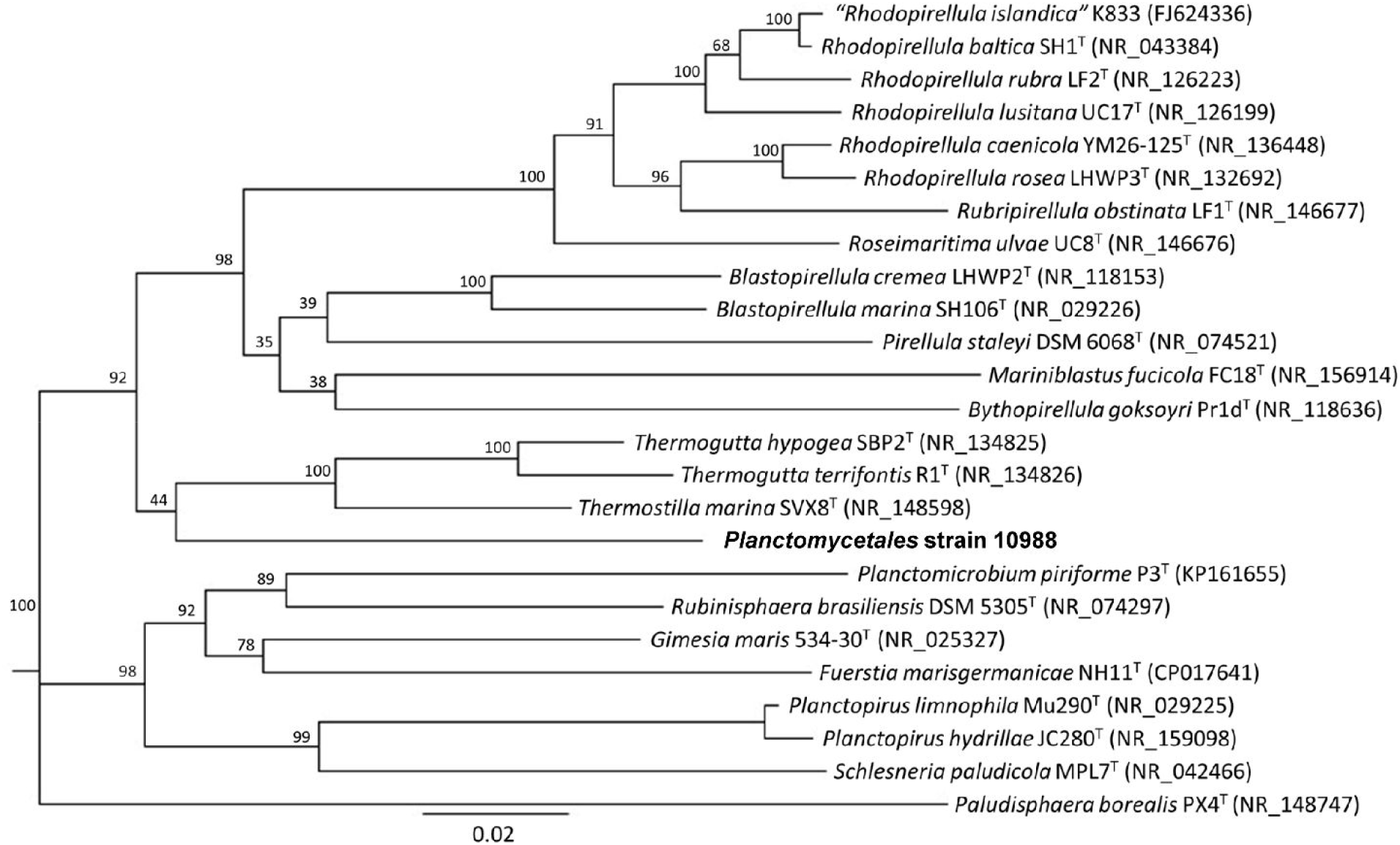
*Phylogenetic tree inferred from 16S rRNA gene sequence similarity showing* Planctomycetales *strain 10988 among its nearest neighbors in the family* Planctomycetaceae. *GenBank accession numbers are indicated in parenthesis.* Paludisphaera borealis *strain PX4^T^ was used as outgroup to root the tree. The scale bar indicates nucleotide substitutions per site.*

The nearest neighbors of strain 10988 among the *Planctomycetales* are the genera *Thermogutta* and *Thermostilla*, both of which are thermophilic as well as anaerobic or microaerophilic. However, strain 10988 showed very different characteristics. It does not only display optimum growth at 24 to 37 °C, but it also grows aerobically. These findings agree with our 16S rDNA gene classification attempt, which classified strain 10988 as only distantly related to all characterized *Planctomycetes*. We thus sought to evaluate its secondary metabolome as a potential source for new natural products. Although *Planctomycetes* have been continuously studied since the late 1980s and *Planctomycetes* have shown to possess cellular features completely distinct from other prokaryotes, little has been done to investigate the planctomycetal secondary metabolome.^15,16^ One of the key limiting factors that need to be overcome in order to uncover planctomycetal secondary metabolism is the requirement to develop suitable cultivation techniques first. While there has been some success with cultivation of freshwater *Planctomycetes*, cultivation of marine species such as strain 10988 turned out to be challenging.^17^ Cultivation of strain 10988 showed that it is an obligate halophile, as it did not grow in absence of sea salts. The halophilicity of the strain is underpinned by an ectoine biosynthesis gene cluster present in genome of strain 10988 that serves as a means to counterbalance the osmotic stress exerted by sea water brine on the cell. Furthermore, the strain depends on surface adsorption for efficient growth which is exemplified by its enhanced growth in early stages if cellulose powders are added to the medium. As these fine filter paper pieces turn to rose color before the medium in the shake flask contains a significant number of suspended cells, the bacterium seems to preferentially colonize surfaces before dispersing into suspension in a shake flask (supporting information).

This finding is well in line with planctomycetal growth in nature that occurs in parts fixed to the surfaces of macroalgae.^9^ The slow growth of this isolate in combination with initially low secondary metabolite production rates – as judged by LC-MS analysis - led us to devise a fermenter based cultivation to obtain increased secondary metabolite yields that could not be achieved in shake flask cultivations. As a means to stimulate productivity of the planctomycetal strain for secondary metabolite isolation, we added adsorber resin to shake flask cultures. This should circumvent productivity limitations arising from feedback inhibition mechanisms.^18^ However, addition of adsorber resin led to complete suppression of planctomycetal growth unless the culture was inoculated with a high concentration of actively growing cells. When strain 10988 was grown in absence of adsorber resin, inoculation of liquid cultures with a very low concentration of live cells was sufficient to stimulate planctomycetal growth. The most probable explanation for this phenomenon is that the presence of adsorber resin in low density cultures masks certain quorum sensing signals by binding them, inhibiting cooperative growth of planctomycetes. While quorum sensing has been linked to different effects such as the inhibition of biofilm formation or virulence, a quorum sensing signal that increases or stalls cell division speed has not been described yet.^19^ As a result, in order to avoid lack of growth or unnecessarily lengthy lag phases in planctomycetal fermentations in larger scale production cultures, adsorber resin addition was performed several days post inoculation of the respective fermenter. In analytical scale shake flask cultivations, the effect of adding resin directly could be mitigated by inoculation of the cultures with a higher concentration of live cells.

### 2.2 Discovery of 3,5-dibromo p-anisic acid

To assess the secondary metabolome of planctomycetal strain 10988, methanolic extracts of the strain’s culture supplemented with adsorber resin were prepared and compared to methanolic extracts of the corresponding medium (“blank” sample) to obtain an overview about its secondary metabolome. The bacterial extract as well as the blank were subjected to high-resolution LC-MS analysis using a reverse-phase UPLC-coupled qTOF setup (supporting information). This analysis revealed an intriguing signal presenting a monoisotopic mass of 308.873 Da [M+H]^+^ (C_8_H_6_Br_2_O_3_, Δ m/z = 6 ppm) and an isotope pattern that pointed towards double bromination.^20^ Maximum cell density as well as the production rate of this doubly brominated compound remained limiting for material supply in shake flask cultures even after media optimization (supporting information). We therefore developed a method to grow strain 10988 in a fermenter which allowed to purify the candidate compound by semi-preparative HPLC (supporting information).

**Figure 3.**
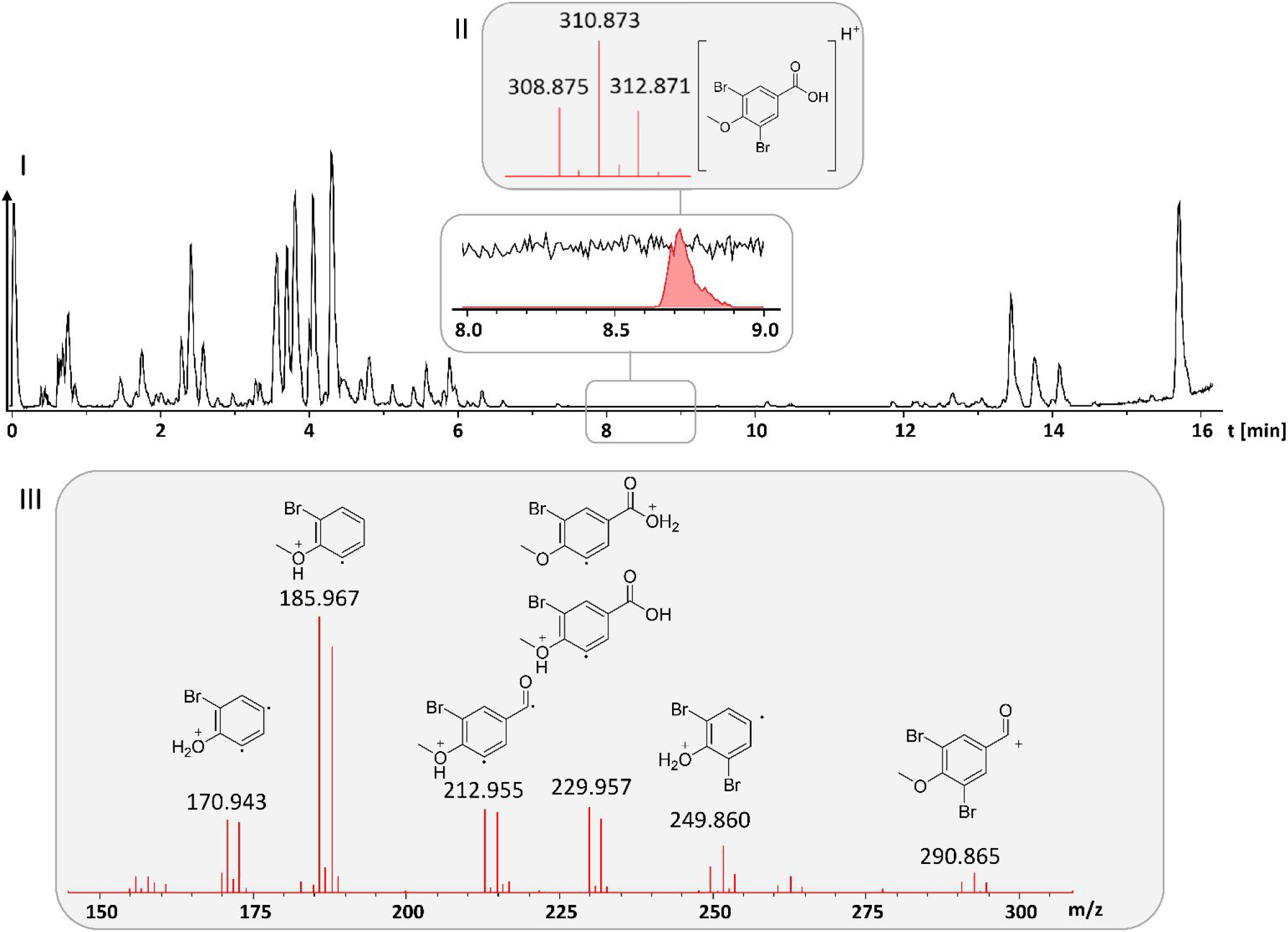
*I) LC-MS chromatogram of* Planctomycetales *strain 10988 with magnification of the MS signal for 3,5 dibromo p-anisic acid (**1**) II) Corresponding MS spectrum and structure formula of **1** III) MS^2^ spectrum of **1** and putative product ions formed in MS^2^ fragmentation.*

According to molecular formula calculation the molecule possesses 6 hydrogen atoms. As the ^1^H NMR spectrum contains only 2 singlet signals, the molecule has to be highly symmetrical. One of the signals representing 2 protons has a strong downfield shift of 8.11 ppm indicating a heavily electron deficient symmetrical aromatic system while the other singlet signal representing 3 protons is characteristic for an oxygen-linked methyl group. The ^13^C shift of the corresponding methyl group indicates its connection to the phenolic oxygen of the molecule and not to the carboxylic acid (supporting information). This is further supported by the fact that the tandem MS spectra show a strong water loss as expected from free carboxylic acid moieties, while we did not observe neutral loss of methanol in tandem MS experiments (Figure 3). This neutral loss would be expected if the molecule contained a methyl ester. The double brominated compound produced by 10988 was therefore determined to be 3,5 dibromo-p-anisic acid (**1**), which could be later confirmed using synthetic standard material.

### 2.3 Biosynthesis of 3,5-dibromo p-anisic acid

In order to identify the biosynthetic origin of **1**, we determined the complete genome of *Planctomycetales* strain 10988 using PacBio long read sequencing technology (supporting information). Genome assembly resulted in a single circular bacterial chromosome of 6.6 Mbp with a total GC content of 50.4% (GenBank accession number XXX). AntiSMASH analysis of the bacterial genome annotated 3 terpene biosynthetic gene clusters (BGCs), an ectoine BGC, a cluster for lassopeptide biosynthesis and a PKS type III BGC.^21^ Contrary to earlier genome mining results from planctomycetes, the genome of strain 10988 does not encode any multimodular secondary metabolite pathways in its genome.^11,22^ Since our newly elucidated secondary metabolite is likely not produced by a multimodular megasynthase, the biosynthesis gene cluster predictors run by antiSMASH are in this case unsuitable to annotate the corresponding biosynthesis pathway, which possibly consists of a set of ‘stand-alone’ enzymes.^21^ We therefore searched the obtained genome for flavin dependent halogenase enzymes, as most region selective bromination or chlorination reactions on aromatic systems are catalyzed by this protein family in nature. ^23,24^ The flavoprotein showing highest homology to halogenating enzymes was named BaaB. It was found encoded adjacent to - and putatively on the same mRNA strand as - a chorismate lyase-like protein termed BaaA. This protein could plausibly deliver the precursor para-hydroxy benzoeic acid from the cellular chorismic acid pool (Figure 4). ^25^ Homology modelling of the two proteins on the protein fold recognition server Phyre2 supports this finding, as both proteins involved in biosynthesis of **1** are correctly mapped onto the expected protein families.^26^ Unfortunately, and despite serious efforts we were unable to develop methods to genetically manipulate the planctomycetal strain and we were thus unable to further validate our hypothesis by an inactivation mutant of the *baaA* and *baaB* locus. The mechanism of UbiC-like chorismate lyases such as BaaA is less studied in comparison to the mechanism of chorismate mutases. Chorismate mutase reactions consist of an electrocyclic 6 electron rearrangement reaction that leads to prephenate formation (Figure 4).^25^ Chorismate lyase enzymes like BaaA use a closely related electrocyclic 6 electron rearrangement reaction that removes pyruvate from chorismate to form p-hydroxy benzoeic acid.^27^ At this point we cannot differentiate whether p-hydroxybenzoeic acid is methylated to p-anisic acid first or if methyl transfer occurs after double bromination of p-hydroxybezoeic acid by the brominase enzyme BaaB. In order to finish biosynthesis of **1** after the action of BaaA and BaaB, an SAM dependent O-methyl transferase (BaaC) is needed that transforms **2** into **1**. When analyzing the genetic locus encoding BaaA and BaaB we did not find such an enzyme, meaning BaaC is encoded in a different genetic locus. The fact that the *baaA* and *baaB* genes are encoded adjacently in the genome, while the corresponding methyl transferase *baaC* is encoded in a different location may indicate that dibromo p-hydroxybenzoeic acid is produced fist and subsequently methylated. It is worth noting that we could not identify either of the possible intermediates *via* LC-MS from the 10988 strain extracts. As BaaB is the only halogenase enzyme encoded in the *baa* BGC, it is certainly responsible for 3,5 dibromination of the aromatic moiety. As both positions that are brominated are chemically equivalent it is not surprising that both halogenations are performed by the same enzyme. Furthermore, we observe strict specificity of BaaB for bromine as no chlorinated or mixed brominated and chlorinated anisic acid derivatives can be identified in the fermentation broth. Thus, BaaB is either unable to bind chloride anions instead of bromide anions due to a difference in binding cavity size, or the redox potential of BaaB is not sufficient to oxidize chloride anions but is sufficient to oxidize bromide ions to an activated species. Still, BaaB is not the only such enzyme unable to process chlorine, as the brominase Bmp5 from *Pseudoalteromonas* strains involved in biosynthesis of polybrominated phenols is also specific for bromine over chlorine.^28^ The architectures of the responsible loci producing polybrominated biphenylic secondary metabolites show remarkable similarity to the Baa operon even though the host organisms are phylogenetically very distant. While Bmp5-like proteins from *P. luteoviolacea* 2ta16 and *P. phenolica* O-BC30 are very similar as they share 96% homology, their similarity to BaaB remains around 44%. This finding is readily explained as *Pseudoalteromonads* and *Planctomycetes* are phylogenetically far apart and BaaB only catalyzes meta-position bromination, while Bmp5 also catalyzes ipso substitution of CO2 at the aromatic core. This reaction removes the carboxylic acid and the phenols are not methylated afterwards.^29,30^ Furthermore, in biosynthesis of polybrominated biphenyl ethers in *Pseudoalteromonads*, an additional enzyme called Bmp7 uses phenolic coupling reactions to form biphenyl structures that do not exist in our planctomyces strain, as the *baa* gene cluster in strain 10988 does not possess the corresponding CYP P450 enzymes.^30^ The absence of CYP enzymes in the planctomycetal BGC explains why the planctomycete only synthesizes monocyclic polybrominated aromatic compounds, as it lacks the CYP enzyme required to perform phenol couplings leading to the formation of biphenylic compounds.^30,31^

**Figure 4.**
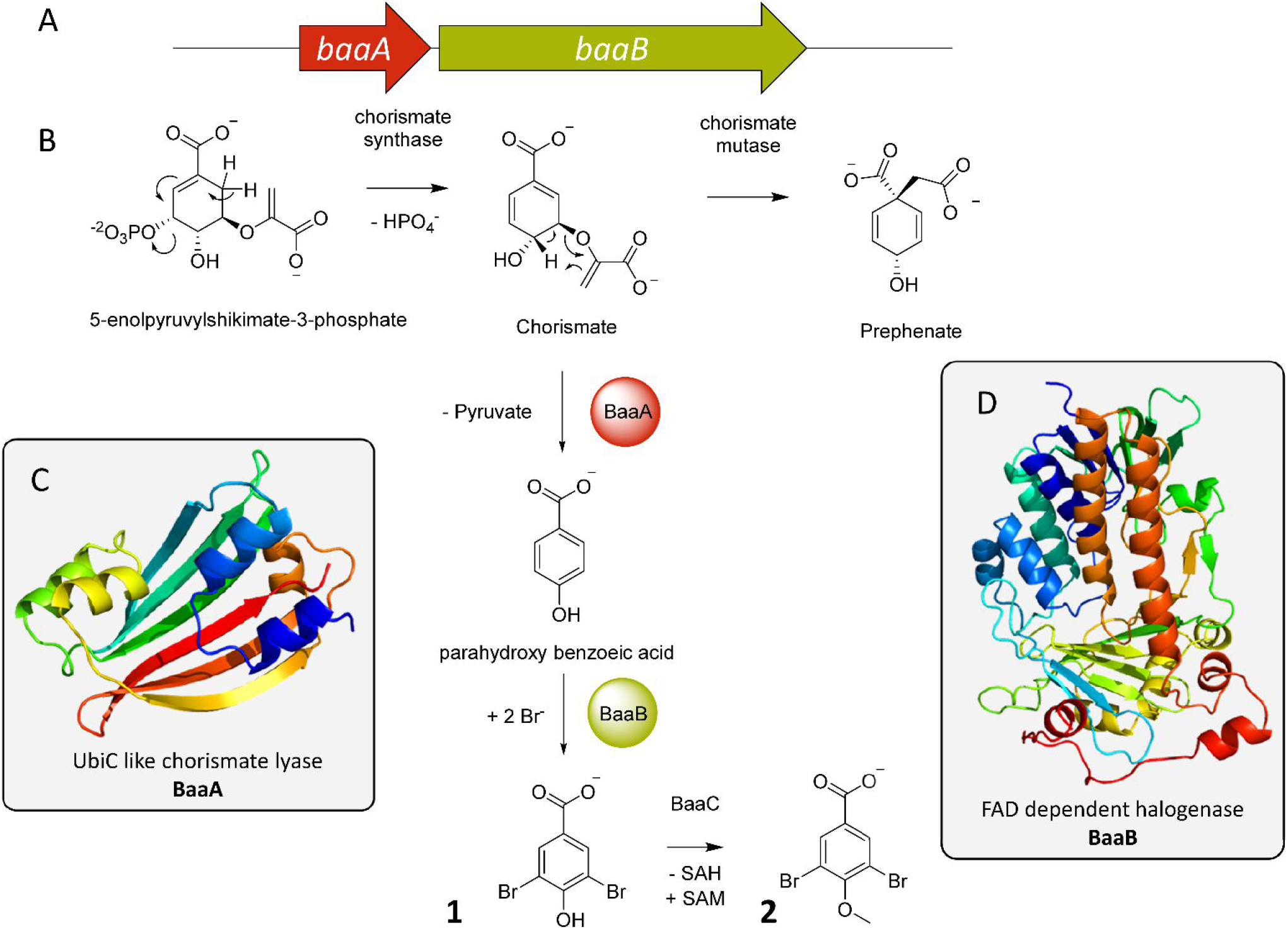
*A) Gene cluster for 3,5 dibromo p-anisic acid (**1**) biosynthesis B) Proposed model for biosynthesis of **1** including the non-methylated precursor **2** C) Phyre2 based homology model for the chorismate lyase BaaA D) Phyre2 based homology model for the brominase BaaB.*

As shown in Figure 4, in order to finish biosynthesis of **1**, the formerly mentioned oxygen-methyl transferase BaaC is required that transfers a methyl group and thus forms the methoxy group in **1**. The corresponding oxygen-methyl transferase responsible for methyl transfer to the phenolic oxygen of the precursor compound is not clustered with the genes responsible for p-anisic acid production. Although there are some candidate sequences for the *BaaC* protein, it is impossible to exactly pinpoint the methyl transferase in the 10988 genome responsible for methylation of **2** due to our inability to perform directed mutagenesis with the strain. Nevertheless, following blast analysis of the 10988 genome for homologues to the StiK protein from stigmatellin biosynthesis in *S. aurantiaca* (NCBI protein acc. Nr. CAD19094.1) and UbiG from *Escherichia coli* K12 (NCBI protein acc. Nr. BAA16049.1) that both catalyze methyl transfer to an aromatic hydroxyl group, we obtained 8 candidate sequences.^32,33^ These candidate genes for the enzyme BaaC, which show similarity to both aforementioned enzymes, can thus be assumed to catalyze reactions such as the transformation of **2** to **1**.

### 2.4 Bioactivity evaluation of 1 and its analogs

Due to the double bromination of **1**, its biosynthesis is costly to the strain in terms of energy and resources. Therefore, **1** can be plausibly expected to confer a competitive advantage to strain 10988 in its environment. To evaluate this biological role of **1** we set out to profile its bioactivity as well as the bioactivity of its biological precursor **2** and its isomer Methyl 3,5 p-hydroxy benzoeic acid (**3**). The compounds **2** and **3** that cannot be obtained from the planctomycetal culture broth are commercially available.

**Scheme 1.**
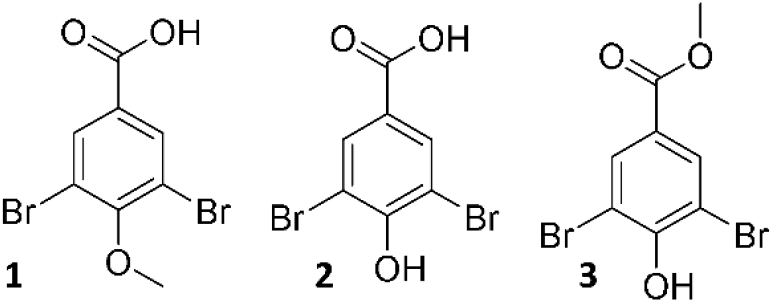
*The natural product 3,5 dibromo p-anisic acid (**1**), its putative biological precursor 3,5 dibromo p-hydroxybenzoeic acid and the natural product analog Methyl-3,5 dibromo p-hydroxybenzoate (**3**)*

The planctomycetal natural product 1, its precursor 2 and its analog 3 were tested in an antibiotics screening against the bacterial pathogens *C. freundii, A. baumanii, E. coli, M. smegmatis, S. aureus, P. aeruginosa, B. subtilis* and *M. luteus*, the yeasts *C. albicans* and *P. anomala* as well as the filamentous fungus *M. hiemalis*. The compounds did not display any inhibition of these microbial indicator strains at concentrations up to 64 μg/ml. To evaluate cytotoxicity we tested **1**, **2** and **3** in a cell line cytotoxicity assay. This assay revealed both methylated compounds (**1** and **3**) to display moderate cytotoxicity to the human cervical carcinoma cell line KB3.1. While **1** and **3** showed an IC50 value of 60 μg/ml, no cytotoxicity was found for the free acid **2** up to 300 μg/ml. To assay herbicidal activity of p-methoxy dibromo benzoeic acid as well as its precursor and isomer we tested the compound’s activity on the germination of *Agrostis stolonifera* penncross. IC50 values for seed germination inhibition were determined to be 32 μg/ml for 1, 64 μg/ml for **2** and 16 μg/ml for **3**. The fully decorated methoxy derivative **1** shows higher biological activity than its precursor **2** while the non-natural methyl ester derivative **3** shows the best anti-germination activity in the *A. stolonifera* germination assay (supporting information). As compound **1** shows no significant antibacterial activity or mammalian cell cytotoxicity but displays moderate phytotoxicity and as **1** is likely released to the marine environment, we assume its role as a putative algal toxin (supporting information).

### 2.5 On the potential biological role of 1

Many planctomycetes live in close association with macroalgae which they cover almost completely.^34^ Given that the strain 10988 was isolated from a marine sediment, the strain is likely associated with marine algae in its natural habitat. The exact mode of interaction between the planctomycetes and the macroalgae has yet to be determined. Still, the high abundance of planctomycetes on algal species, which can reach up to 50 % of the algal microbiome, indicates that these bacteria are significant interaction partners for the algae.^34^ One hypothesis considers the algae as a food source for planctomycetes, since they are able to utilize uncommon sugars such as rhamnose and fucose contained in algal biomass. In our case, strain 10988 – like other planctomycetes – was able to grow on uncommon sugars such as galactose, mannose, lactose, sucrose, maltose raffinose, xylose and rhamnose (supporting information, Figure S2), indicating that this nutritional option could apply to strain 10988.^11^ As on the other hand, the planctomyces bacterium possesses the ability to produce the plant toxin 3,5 dibromo p-anisic acid, whose production seems to be tightly controlled as judged by the low production titers under laboratory conditions, we reason that an ambivalent interaction model might take place between planctomycetes and their plant hosts. The planctomycetes strains probably live on the algal surface and modulate the local microbial community until they sense the algal species they live on is weakening. This might trigger expression of said plant toxin to kill and decay this part of the algae and the bacterium would subsequently move on to colonize different algae. Similar ‘Jeckyll and Hyde’ behavior, meaning the ability to switch between commensalism and symbiosis, comparable to planctomycetal colonization of algae, and a virulent state that is hostile to its host organism has been described for the human pathogen *C. albicans*.^35^ The ability of strain 10988 to produce a plant toxin as a bacterium associated to macroscopic plants can be seen as a strong hint that the bacterium adopts such a strategy.

## 3 Conclusion

In this work we describe the cultivation of a new marine planctomycete that is genetically distant from all planctomycetes known to date, and reveal *Planctomycetales* strain 10988 as producer of a dibrominated secondary metabolite. Isolation of this secondary metabolite required a stirred tank reactor setting and optimized medium and culture conditions. Subsequent structure elucidation of **1** by NMR revealed an intriguing structure and thus sparked interest in the biosynthetic origin and ecological role of the compound. We were able to pinpoint the core biosynthesis genes *baaA* and *baaB* that can accomplish the core structure of **1**. Investigation of the bioactivity of **1** as well as the bioactivity of its isomer **3** and putative precursor **2** showed that this compound class displays herbicidal activity in *A. stolonifera* penncross germination assays, leading us to hypothesize on a biological role of this compound in the life cycle of the algal symbiont *Planctomycetales* strain 10988. In conclusion, we contribute to the understanding of the biogenesis of small polyhalogenated compounds in marine bacteria, whereas it remains astonishing to what extent such polybrominated aromatic substances are apparently released into the ecosystem from biological instead of anthropogenic sources.

This study also identifies planctomycetes as an underexploited source of biologically active secondary metabolites, as **1** to the best of our knowledge is the first natural product described from this bacterial taxon. However, we would like to point out that previous studies may have overestimated the genome encoded secondary metabolite diversity of planctomycetes as a group, since strain 10988 under study here did not show the presence of any multimodular megasynthetase.^11,22^ Even though strain 10988 shows some BGCs, especially BGCs linked to terpene biosynthesis, megasynthase containing BGCs are often considered as benchmark indicators for secondary metabolite production capability.^2^ On the other hand, the example presented here shows how important it is to evaluate new taxa on the metabolomics stage, since metabolites of the type described here are easily missed by genome mining. Thus, the overall potential of planctomycetes awaits further investigation. While growing these bacteria under laboratory conditions may be tedious and non-trivial, devising methods for their cultivation is a valuable tool to tap into the planctomycetal secondary metabolite space. The discovery of polybrominated compounds in strain 10988 is well in line with both the observation that this bacterium is an obligate halophile and reminiscent of the strain’s marine origin. The isolated and characterized natural product 3,5 dibromo p-anisic acid shows that Nature, especially the marine microbial community is able to biosynthesize polyhalogenated small aromatic compounds that look like anthropogenic products of chemical synthesis.

## Acknowledgement

The authors would like to thank Anja Paluczak and Viktoria Schmitt for the herbicidal and antimicrobial assays with 3,5 dibromo p-anisic acid and its analogs. We thank Cathrin Spröer for genome sequencing and Simone Severitt and Nicole Heyer for excellent technical assistance.

## Supporting Information

An in detail description of the planctomyces strain, all utilized fermentation protocols, *in silico* analyzes on gene and protein level as well as all relevant NMR data for structure elucidation are available free of charge via the Internet at BioRxiv.org.

