## Supporting information for "Discovery of a polybrominated aromatic secondary metabolite from a planctomycete points at an ambivalent interaction with its macroalgae host"

### 1 Growth of *Planctomycetales* strain 10988

#### 1.1 Isolation and enrichment of *Planctomycetales* strain 10988

*Planctomycetales* strain 10988 was isolated from sand sample collected in Batangas, Philippines using an isolation set-up for marine myxobacteria. The sand sample was placed on an agar plate [Ferric citrate 0.01 g/L, $MgSO_4 \cdot 7H_2O$ 8.0 g/L, $CaCl_2 \cdot 2H_2O$ 1.0 g/L, NaCl 3 g/L, KCl 0.5 g/L, $NaHCO_3$ 0.16 g/L, $H_3BO_3$ 0.02 g/L, KBr 0.08 g/L, $SrCl_2 \cdot 6H_2O$ 0.03 g/L, di-Na-ß-glycerophosphate 0.01 g/L, trace element solution 1.0 mL/L, BD Bacto Agar 15 g/L, HEPES 1.19 g/L, dissolved in DI water, pH adjusted to 7.2 with NaOH before autoclaving] and baited with 4 strips (1 x 1cm) of sterile filter paper. The strain was detected by its ability to form a thin and transparent swarming colony radiating from the inoculum on the agar plate. After repeated cutting of the swarm edge and transferring it to the agar same medium supplemented with autoclaved Baker's yeast (5 g/L), the strain was isolated and purified. The strain can be stored as a 50% glycerol stock remaining viable at -80 °C.

#### 1.2 Media recipes used in planctomycetes cultivation

*Planctomycetales* strain 10988 was grown in solid and liquid culture on PYMA medium to propagate the strain. For creation of solid media 16 g/L agarose (Sigma Aldrich) are added to the liquid medium before autoclaving.

*Table 1. Medium recipe of PYMA medium used to propagate 10988*

| PYMA Medium | | |
|---|---|---|
| Ingredient | Concentration | Supplier |
| Sea Salts | 36 g/L | ATI |
| trizma base | 10 mM | Sigma Aldrich |
| Peptone from soy meal | 0.15 g/L | Merck |
| yeast extracs | 0.15 g/L | BD |
| D(+) maltose monohydrate | 1 g/L | Sigma Aldrich |
| $NH_4SO_4$ | 0.25 g/L | VWR |
| pH is adjusted to 7.5 with HCl before autoclaving | | |

For fermentation, the medium was optimized to produce a maximal amount of 3,5 dibromo p-anisic acid. As yield evaluations point to ECX medium as the best tested medium in terms of 3,5 dibromo p-anisic acid production fermentative production was performed in ECX medium.

*Table 2. Medium recipe for ECX medium used for fermentative production of 3,5 dibromo p-anisic acid*

| PYMA Medium | | |
|---|---|---|
| Ingredient | Concentration | Supplier |
| Sea Salts | 32 g/L | ATI |
| NaBr | 4 g/L | Sigma Aldrich |
| trizma base | 10 mM | Sigma Aldrich |
| Peptone from soy meal | 2 g/L | Merck |
| Meat extract | 1 g/L | Merck |
| Lactose | 1 g/L | Sigma aldrich |
| Sucrose | 1 g/L | Roth |
| Xylose | 1 g/L | ACROS |
| Raffinose | 1 g/L | TCI biochemicals |
| Casitone | 1 g/L | BD |
| D(+) maltose monohydrate | 4 g/L | Sigma Aldrich |
| Cellulose fiber (Grade BER 40) | 2 g/L | Access Bio |
| pH is adjusted to 7.5 with HCl before autoclaving, add 200 µg/L Vitamin B12 solution after autoclaving | | |

#### 1.3 Shake flask cultivation of strain 10988

Cultures for UHPLC/HRMS analysis are grown in 300 ml shake flasks containing 50 ml of ECX or PYMA medium for strain 10988 inoculated with 1 ml of pre-culture. After inoculation the medium is supplemented with 2% (v/v) of sterile XAD-16 adsorber resin (Sigma Aldrich) suspension in water if the culture is to be extracted. For propagation of the cells in liquid medium, the cultures are grown without XAD-16 adsorber resin. Strain 10988 is cultivated at 28°C for 21 days on an orbitron rotatory shaker at 160 rpm.

#### 1.4 Investigation of growth and production kinetics of *Planctomycetales* strain 10988

In order to elucidate at what growth stage 10988 produces our target compound we chose to investigate secondary metabolite growth kinetics and production kinetics of 3,5 dibromo p-anisic acid at the same time. 18 separate sets of cultures are grown according to the shake flask growth protocol, which are all inoculated by the same pre culture. Every two days, two shake flasks are harvested and extracted according to the analytical scale extraction protocol. Before centrifuging the culture, 1 ml of culture broth is taken out after the cellulose in the medium has settled in order to measure the cultures OD 600 value that is proportional to cell density with an Eppendorf BioPhotometer plus. If the OD 600 approaches 0.5 or higher, the OD is measured from serial dilutions to ensure accuracy of the value. Purity of the strain is checked under a light microscope to ensure measuring OD 600 values from contamination free cultures. The rest of the culture is extracted according to the analytical scale extraction protocol for UHPLC-MS analyses and analyzed via UHPLC-MS according to standard conditions. 3,5 dibromo p-anisic acid is quantified by peak area of the most intense isotopic ion of 310.873 Da $[M+H]^+$.

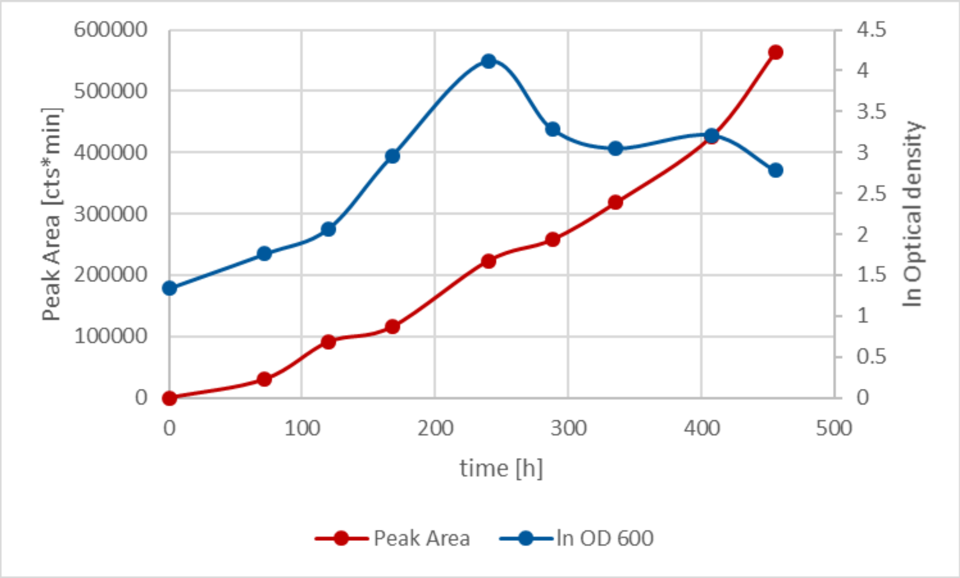

*Figure 1. Growth and production kinetics of 3,5 dibromo p-anisic acid in 10988*

It should be noted that after approximately 11 days the cell density peaks. Afterwards, dying cells start forming cell clumps and debris, which makes the optical density fall faster than one would expect if dead cells would be kept in suspension.

*Table 3. Optical density of strain 10988 cultures and peak area of 3,5 dibromo p-anisic acid in the culture extract*

| t [h] | OD 600 Shake flask 1 | OD 600 Shake flask 2 | Peak area Shake flask 1 | Peak area Shake flask 2 |
|---|---|---|---|---|
| 0 | 2,61 | 1,21 | 0 | 0 |
| 72 | 2,71 | 3,1 | 25503 | 35219 |
| 120 | 4,44 | 3,48 | 77993 | 104681 |
| 168 | 8,3 | 11,0 | 96643 | 135500 |
| 240 | 37,1 | 24,0 | 270906 | 176006 |
| 288 | 13,2 | 13,6 | 212728 | 302992 |

| | | | | |
|---|---|---|---|---|
| 336 | 11,9 | 9,2 | 290473 | 345969 |
| 408 | 17,2 | 7,6 | 453838 | 398107 |
| 456 | 7,2 | 8,9 | 623366 | 503019 |

### 1.5 Cultivation of strain 10988 in batch bioreactors

#### 1.5.1 Fermentation optimization in 1L bioreactors

In order to optimize fermentation for efficient production of 3,5 dibromo p-anisic acid we tested the parameters in an 1L Infors HT fermenter (Art. No. 26133) equipped with two 6 blade plate stirrers with a diameter of 5cm. The fermenter is gassed with synthetic air (1.5 bar), $pO_2$ and pH are logged with respective electrodes during fermentation. The fermenter is connected to 4 different Schott bottles containing 1M $H_2SO_4$, 1M NaOH for pH control, an 1/10 dilution of Antifoam Y-30 Emulsion (Sigma) in DI water and an empty Schott bottle for inoculation. The bioreactor is filled with 1L of ECX medium supplemented with 5 ml of undiluted Antifoam Y-30 Emulsion. After autoclaving at 121°C for 30 min, electrodes are connected to the fermenter and the default values for the fermentation are put in. pH is set to 7.2, stirring speed to 150 rpm and fermentation temperature is set to 28°C. Temperature is taken from the fermentation broth via a thermocouple that is planted in a blind tube in the fermentation broth. Inoculation is done with an autoclaved glass syringe and a steel needle that can be flame sterilized before inoculation. The fermenter is inoculated with 50 ml of a well grown planctomycetal pre culture in a 300 ml shake flask filled with 50 ml ECX medium. The OD600 value should exceed 2 and the culture should be intensely rose colored. This inoculation broth is injected into the empty schott bottle and introduced into the fermenter via excess air pressure. After 5 to 7 days depending on the growth speed, 20 ml of a sterile XAD-16 adsorber resin solution (Sigma Aldrich, 50/50 vol %) are added to the fermenter to remove excess 3,5 dibromo p-anisic acid from the broth and prevent feedback inhibition. Fermentation is done after 15 to 21 days depending on the lag phase. Fermentation broth is harvested by centrifugation at 16000 x g for 10 min with a Beckman Coulter Avanti J-26XP Centrifuge and the supernatant is discarded.

#### 1.5.2 Fermentation in 5L bioreactors

Fermentation is performed in a 5L Infors HT bioreactor (Art. No. 26128) equipped with two 6 blade plate stirrers with a diameter of 5cm. Fermentation is performed largely analogous to the 1L bioreactor fermentation experiments. Inoculation volume is changed to 200 ml pre culture broth to ensure good colonization of the fermenter and stirring speed is elevated to 200 rpm. As for the 1L bioreactor experiments, 200ml sterile XAD-16 adsorber resin solution (Sigma Aldrich, 50/50 vol %) is added to the fermenter broth after 7 days of fermentation. Fermentation end point is determined after 16 to 20 days depending on the batch and workup is performed analogous to the 1L fermentation.

### 2 Phenotypical characterization

The *Planctomycetales* stain 10988 occurs in three distinct phenotypes on agar plates. The strain occurs as a fast swarming white or rose color phenotype and a slower swarming green phenotype. In liquid medium, the strain showed uniquely the rose color phenotype

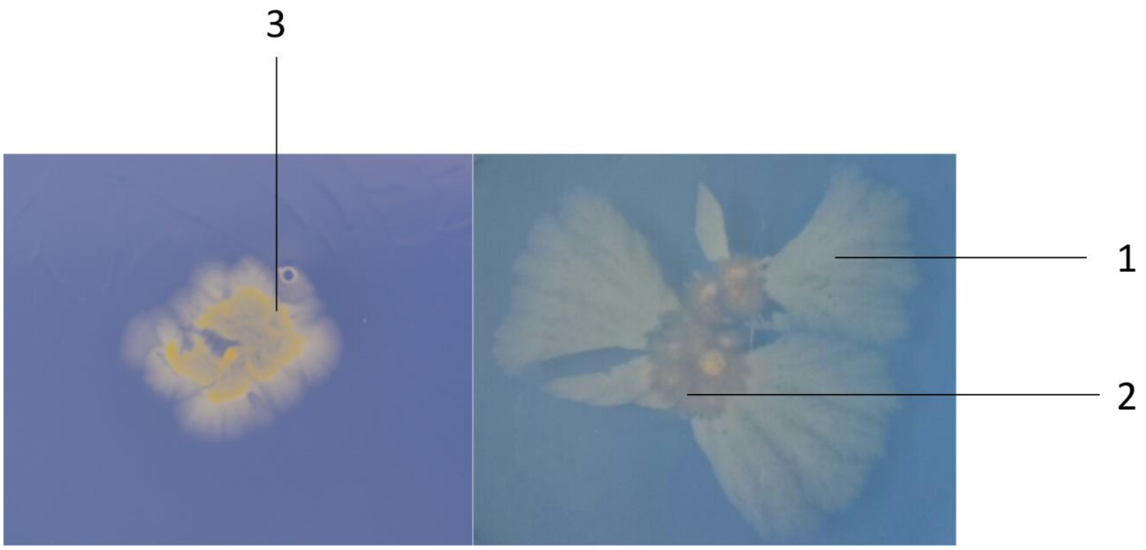

*Figure S 1. Images of the three distinct phenotypes of 10988 in PYMA agar depicting the white (1), rose (2) and green phenotype (3)*

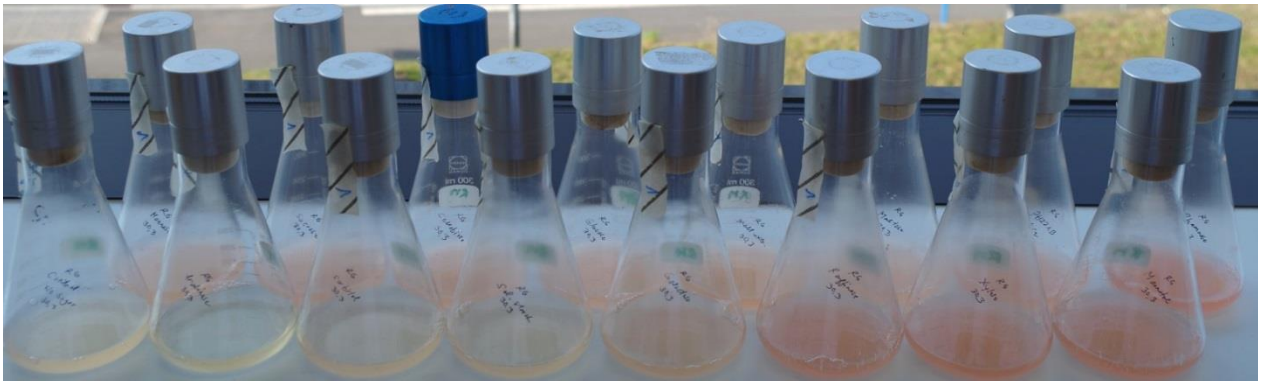

*Figure S 2. Image of the rose color phenotype observed in liquid suspension culture supplemented with different sugar sources.*

### 3 Analytical setting for strain 10988 metabolomics

#### 3.1 Extraction of analytical scale cultures for UPLC-MS analysis

The frozen cell pellet is transferred into a 100 ml Erlenmeyer flask and a magnetic stirrer is added. 50 ml of methanol (fluka, technical grade) are added onto the pellet and the mixture is stirred for 60 min. on a stirring plate. The extract is left to settle in order to sediment cell debris and XAD resin for a second extraction step. The supernatant is filtered through a 125 micron folded filter keeping cell pellet and XAD-16 resin in the Erlenmeyer flask for a second extraction step. The residual pellet and XAD-16 resin is extracted again with 30 ml of acetone (fluka, technical grade) for 60 min on a stirring plate and filtered through the same folded filter. The combined extracts are transferred into a 100 ml round bottom flask. The solvent is evaporated on a rotary evaporator at 260 mbar and 40 °C water bath temperature. The residual water is evaporated at 20 mbar until the residue in the flask is completely dry. The residue is taken up in 500 µl of methanol (Chromasolv HPLC grade, Sigma Aldrich) and transferred into a 1.5 ml Eppendorf tube. This tube is centrifuged with a table centrifuge at 21500 rcf for 5 minutes to remove residual insolubilities such as salts, cell debris and XAD fragments.

#### 3.2 Standardized UPLC-MS conditions

All measurements were performed on a Thermo Scientific (Bremen, Germany) Ultimate 3000 RSLC system using a Waters (Eschborn, Germany) BEH C18 column (50 x 2.1 mm, 1.7 µm) equipped with a Waters VanGuard BEH C18 1.7 µm guard column. Separation of 1 µl sample volume was achieved by a linear gradient from (A) $H_2O$ + 0.1 % FA to (B) ACN + 0.1 % FA at a flow rate of 600 µL/min and a column oven temperature of 45 °C. Gradient conditions were as follows: 0 – 0.5 min, 5% B; 0.5 – 18.5 min, 5 – 95% B; 18.5 – 20.5 min, 95% B; 20.5 – 21 min, 95 – 5% B; 21-22.5 min, 5% B. UV spectra were recorded by a DAD in the range from 200 to 600 nm. The LC flow was split to 75 µL/min before entering the Bruker Daltonics maXis 4G hrToF mass spectrometer (Bremen, Germany) using the Apollo II ESI source. Mass spectra were acquired in centroid mode ranging from 150 – 2500 m/z at a 2 Hz full scan rate. Mass spectrometry source parameters are set to 500V as end plate offset; 4000V as capillary voltage; nebulizer gas pressure 14.5 psi; dry gas flow of 5 l/min and a dry temperature of 200°C. Ion transfer and quadrupole settings are set to Funnel RF 350 Vpp; Multipole RF 400 Vpp as transfer settings and Ion energy of 5eV as well as a low mass cut of 300 m/z as Quadrupole settings. Collision cell is set to 5.0 eV and pre pulse storage time is set to 5 µs. Spectra acquisition rate is set to 2 Hz. Calibration is done automatically before every UPLC-hrMS run by injection of a sodium formate calibrant solution and calibration on the sodium formate clusters forming in the ESI source. All MS analyses are acquired in the presence of the lock masses $C_{12}H_{19}F_{12}N_3O_6P_3$; $C_{18}H_{19}O_6N_3P_3F_2$ and $C_{24}H_{19}F_{36}N_3O_6P_3$, which generate the $[M+H]^+$ ions of 622.028960; 922.009798 and 1221.990638.

### 4 Isolation, purification and characterization of 3,5 dibromo p-anisic acid

#### 4.1 Isolation of 3,5 dibromo p-anisic acid

The crude fermenter broth consisting of cells and XAD-16 resin are harvested by centrifugation on a Beckmann Avanti J-26 XP with the JLA 8.1 rotor at 6000 rcf. Combined resin and cells are extracted successively with 2 x 250 ml of technical grade methanol (Fluka) and 2 x 250 ml of technical grade acetone (Fluka). The extracts are combined, filtered through a 2 µm pore diameter paper filter and all solvent is evaporated on a rotary evaporator. The crude extract is partitioned between hexane and methanol in a separatory funnel, the methanol phase is dried on a rotary evaporator and the residue is partitioned between ethyl acetate and water. 3,5 dibromo p-anisic acid remains almost exclusively in the ethyl acetate layer. The ethyl acetate layer is dried again and taken in a minimum volume of methanol for flash chromatography. Flash chromatography was performed on a Biotage Isolera one system on a 25 g cartridge filled with C-18 coated silica gel with a column volume of 45 ml. Separation is performed with Water as A and Acetonitrile as B and a flow rate of 50 ml/min. Chromatographic separation contains 2 CV of 95 % A followed by a gradient to 100 % B during 6 CV. The chromatographic run is separated into fractions of 8 ml and those containing 3,5 dibromo p-anisic acid are combined for further purification. Further purification is done using a Dionex Ultimate 3000 SDLC low pressure gradient system on a Phenomenex Luna C18-2 250x10mm 5µm column with the eluents H2O + 0.1% FA as A and ACN + 0.1% FA as B, a flow rate of 5 ml/min and a column thermostated at 30°C. B content is kept at 10 % for 2 minutes followed by a linear gradient to 45 % B during 36 minutes. B content is then ramped to 95 % in 2 minutes and kept at 95 % for 3 minutes to flush the column. B content is ramped back to initial conditions during 30 s and the column is reequilibrated for 1.5 minutes. The compound can be detected with a UV detector at 220 nm wavelength. After drying, 3,5 dibromo p-anisic acid is obtained as a white amorphous solid.

#### 4.2 NMR Data of 3,5 dibromo p-anisic acid

1D and 2D NMR data used for structure elucidation of 3,5 dibromo p-anisic acid is acquired in Methanol-$d_4$ and DMSO-$d_6$ on a Bruker Avance III HD 500 UltraShield spectrometer with a 5 mm TXI cryoprobe ($^1$H at 500 MHz, $^{13}$C at 125 MHz) NMR spectrometer. To elucidate the planar structure of the molecules $^1$H, $^{13}$C as well as H-H DQFCOSY, HSQC and HMBC data are acquired. DQFCOSY, HSQC, HMBC, and ROESY experiments were recorded using standard pulse programs. The NMR signals are grouped in the tables below and correspond to the numbering in the scheme above.

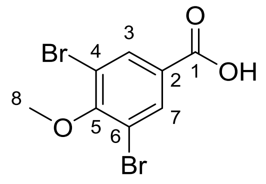

*Figure 2. Structure formula of 3,5 dibromo p-anisic acid including carbon numbering used in NMR analysis*

*Table 4. Tabulated 1D and 2D NMR signal for 3,5 dibromo p-anisic acid in DMSO-$d_6$*

| Carbon number | $^{13}C$ shift | $^{1}H$ shift | J, multiplicity, number of protons | H – H COSY correlations | HMBC correlations |
|---|---|---|---|---|---|
| 1 | 171.3 | - | | - | - |
| 2 | 137.9 | - | | - | - |
| 3,7 | 134.9 | 8.11 | s, 2H | - | 118.1, 134.9, 137.9, 156.8, 171.3 |
| 4,6 | 118.1 | - | | - | - |
| 5 | 156.8 | - | | - | - |
| 8 | 61.0 | 3.87 | s, 3H | - | 156.8 |

*Table 5. Tabulated 1D and 2D NMR signal for 3,5 dibromo p-anisic acid in MeOH-$d_4$*

| Carbon number | $^{13}C$ | $^{1}H$ shift | J, multiplicity, number of protons | H – H COSY correlations | HMBC correlations |
|---|---|---|---|---|---|
| 1 | 164.4 | - | | - | - |
| 2 | 140.1 | - | | - | - |
| 3,7 | 132.6 | 7.95 | s, 2H | - | 116.0, 132.6, 140.1, 152.9, 164.4 |
| 4,6 | 116.0 | - | | - | - |
| 5 | 152.9 | - | | - | - |
| 8 | 60.2 | 3.78 | s, 3H | - | 152.9 |

### 5 Genetic characterization of the Baa gene cluster

#### 5.1 Antibiotics resistance tests for strain 10988

In order to attempt the creation of mutant clones of 10988 to prove the origin of 3,5 dibromo p-anisic acid experimentally we explored the possibility to use resistance markers in 10988. Although there have been some reports on mutagenesis of planctomycetes, only a couple of planctomycetes have been genetically manipulated successfully so far. [4, 6]

*Table 6. Antibiotics sensitivity tests performed on 10988*

| Antibiotic | highest concentration tested [ng/ml] |
|---|---|
| oxytetracycline | 50 |
| kanamycin | 100 |
| ampicillin | 200 |

| | |
|---|---|
| streptomycin | 200 |
| chloramphenicol | 25 |
| apramycin | 50 |
| hygromycin E | 50 |

Unfortunately, the strain showed high resistance to all tested antibiotics. We were therefore unable to perform genetic manipulations with the strain as we could not obtain a viable resistance marker. We were thus also unable to inactivate the 3,5 dibromo p-anisic acid via single crossover homologous recombination.

#### 5.2 Genome sequencing and gene cluster annotation

Genomic DNA of strain 10988 was isolated using phenol/chloroform based Isolation of bacterial genomic DNA. SMRTbell™ template library was prepared according to the instructions from PacificBiosciences, Menlo Park, CA, USA, following the Procedure & Checklist Greater than 10 kb Template Preparation and Sequencing. Briefly, for preparation of 10kb libraries ~10µg genomic DNA was end-repaired and ligated overnight to hairpin adapters applying components from the DNA/Polymerase Binding Kit P4 from Pacific Biosciences, Menlo Park, CA, USA. Reactions were carried out according to the manufacturer´s instructions. SMRTbell™ template was Exonuclease treated for removal of incompletely formed reaction products. Conditions for annealing of sequencing primers and binding of polymerase to purified SMRTbell™ template were assessed with the Calculator in RS Remote, PacificBiosciences, Menlo Park, CA, USA. SMRT sequencing was carried out on the PacBio RS (PacificBiosciences, Menlo Park, CA, USA) taking one 240-minutes movie. PacBio sequencing resulted in 55,213 reads with a mean read length of 12,221 bp. Genome assembly was performed applying the RS_HGAP_Assembly.3 protocol included in SMRT Portal version 2.3.0 applying default parameters and resulted in a one contig sequence covered 85x. The genome of strain 10988 consists of a single circular bacterial chromosome with 6,649,439 base pairs. This gene cluster prediction does not include the genomic locus responsible for the production of 3,5 dibromo p-anisic acid as this polybrominated compound is not biosynthesized by a megasynthase detectable by antiSMASH through hidden markov models. [1] The genome of planctomycetal strain 10988 was deposited at GenBank and can be retrieved under Accession number XXXX.

#### 5.3 Phyre 2 results for modelling the planctomycetal enzymes BaaA and BaaB

When modelling the BaaA protein on the protein fold recognition server Phyre 2 two hit fold structures are retrieved that show a 100 % confidence model based on a PDB structure. The first of these hits was a chorismate lyase from *Archaeoglobus fulgidus* (PDB 2NWI) (Bonanno et al. unpublished results), while the second hit is an *E. coli* chorismate lyase (PDB 1TT8).[5]

Modelling of the BaaB protein on the protein fold recognition server Phyre 2 we could retrieve 60 homology models to PDB crystal structures that show 100 % confidence. All of these proteins folds belong to the class of FAD dependent halogenases and oxygenases. The closest relative of BaaB in the PDB is the halogenase CmlS from the chloramphenicol biosynthesis in *S. venezuelae* (PDB 3I3L) and the second closest relative is the pyoluteorin chlorinating enzyme PltA from *P. aeruginosa* (PDB 5DBJ). [2, 3] Based on these two enzymes BaaA and BaaB being clustered together it is very likely that they are responsible for formation of 3,5 dibromo p-anisic acid.

#### 5.4 Comparison of the BaaB brominase to the Bmp5 enzymes

To assess the degree of structural novelty of the BaaB brominase when compared to their bromophenol producing congeners from Pseudoaltheromonas strains called Bmp5 proteins we compared the brominases on a sequence level. While the two brominating enzymes from *P. luteoviolacea* and *P. phenolica* that have been shown to polybrominate phenols are strikingly similar with a homology of 96 % the homology of the planctomycetal BaaB to either of the Bmp5 proteins is only about 44 %.

*Table S 1. Pairwise homology of BaaB and the Bmp5 enzymes*

| | BaaB<br>Planctomycetal species 10988 | Bmp5<br>*P. luteoviolacea* 2ta16 | Bmp5<br>*P. phenolica* O-BC30 |
|---|---|---|---|
| BaaB<br>Planctomycetal species 10988 | | 43.73 % | 43.96 % |
| Bmp5<br>*P. luteoviolacea* 2ta16 | 43.73 % | | 96.98 % |
| Bmp5<br>*P. phenolica* O-BC30 | 43.96 % | 96.98 % | |

#### 5.5 Candidate enzymes for the O-methyl transferase BaaC

Unfortunately, it became clear that the biosynthetic gene cluster responsible for the production of 3,5 dibromo p-anisic acid does not contain a SAM dependent O-methyl transferase domain protein. When analyzing the Planctomycetes sp. 10988 species by blast using two phenolic O-methyl transferases as probe sequences, we retrieve 8 target sequences that show high similarity to both query sequences. The two query sequences are the O-methyl transferase involved in retrieving these target sequences are the O-methyl transferase StiK responsible for SAM-dependent methyl transfer in Stigmatellin biosynthesis in *S. aurantiaca* (NCBI protein acc. Nr. CAD19094.1) and the *E. coli* K-12 protein responsible for O-methyl transfer in Ubiquinone biosynthesis UbiG (NCBI protein acc. Nr. BAA16049.1) that both catalyze similar reactions. Locus tags of these identified BaaC candidates are Methyltransferase 530, Methyltransferase 21270, Methyltransferase 22400, Methyltransferase 23570, Methyltransferase 27160, Methyltransferase 28040, Methyltransferase 32880 and Methyltransferase 36580. As we do not have any means to perform genetic manipulation on 10988 we have no means of identifying the methyl transferase performing this methylation reaction among these 8 candidates.

### 6 Bioactivity assays

#### 6.1 Origin of the assayed compounds

3,5 dibromo p-hydroxybenzoeic acid (**2**) and Methyl 3,5 dibromo p-hydroxybezoeic acid (**3**) were obtained from the commercial chemical supplier Alfa Aesar (Kandel, Germany). Additional 3,5 dibromo p-anisic acid (**1**) was obtained from Fluorochem Ltd. (Derbyshire, UK). Identity of **1** obtained from a commercial source to its natural product counterpart was assayed by NMR analysis, UHPLC retention time and the LC-$MS^2$ spectrum.

#### 6.2 Antimicrobial Assay

All microorganisms were handled according to standard procedures and were obtained from the German Collection of Microorganisms and Cell Cultures (*Deutsche Sammlung für Mikroorganismen und Zellkulturen*, DSMZ) or were part of our internal strain collection. For micro dilution assays, overnight cultures of Gram-positive bacteria in Müller-Hinton broth (0.2 % (w/v) beef infusion, 0.15 % (w/v) corn starch, 1.75 % (w/v) casein peptone; pH 7.4) were diluted in the growth medium to achieve a final inoculum of ca. $10^6$ cfu $ml^{-1}$. Serial dilutions of the brominated benzoeic acid derivatives were prepared from freshly prepared Methanol stocks in sterile 96-well plates. The cell suspension was added and microorganisms were grown on a microplate shaker (750 rpm, 37°C and 16 h). Growth inhibition was assessed by visual inspection and given MIC (minimum inhibitory concentration) values are the lowest concentration of antibiotic at which no visible growth was observed.

#### 6.3 Cytotoxic Activity

Cell lines were obtained from the German Collection of Microorganisms and Cell Cultures (Deutsche Sammlung für Mikroorganismen und Zellkulturen, DSMZ) or were part of our internal collection and were cultured under conditions recommended by the depositor. Cells were seeded at 6 x $10^3$ cells per well of 96-well plates in 180 µl complete medium and treated with the brominated benzoeic acid derivatives in serial dilution after 2 h of equilibration. Each compound was tested in duplicate as well as the internal solvent control. After 5 d incubation, 20 µl of 5 mg $ml^{-1}$ MTT (thiazolyl blue tetrazolium bromide) in PBS was added per well and it was further incubated for 2 h at 37°C. The medium was discarded and cells were washed with 100 µl PBS before adding 100 µl 2-propanol/10 N HCl (250:1) in order to dissolve formazan granules. The absorbance at 570 nm was measured using a microplate reader (Tecan Infinite M200Pro), and cell viability was expressed as percentage relative to the respective methanol control. $IC_{50}$ values were determined by sigmoidal curve fitting.

#### 6.4 Herbicidal activity assays

*A. stolonifera* penncross seeds are grown in plant medium consisting of Serva 47516.03 Murashige & Skoog plant *salts* (2.2 g/L) and Gamborg's B-5 Basal Medium with Minimal Organics (1.6 g/L) in deionized water. The essay is performed in sterile 96 well flat bottom well plates containing ca. 10 seeds, 195 µL of plant medium and 5 µL of the assayed compound per well. Positive controls are performed using solely plant medium as well as plant medium with methanol. The prepared 96 well plates are incubated for 6 days in a Grow light Garden illumination chamber to germinate the *A. stolonifera* seeds. Inhibition is determined by visual inspection of the respective wells. Herbicidal activity is detected by a lack of germination in the respective wells, wells containing more than 5 germinated plant seeds count as not inhibited.

*Table S 2. 96 well plate layout for the herbicidal assay, Cmp.1 is 3,5 dibromo p-hydroxybenzoeic acid methyl ester, Cmp. 2 is 3,5 dibromo p-anisic acid and Cmp. 3 is 3,5 dibromo p-hydroxybenzoeic acid*

| | 1 | 2 | 3 | 4 | 5 | 6 | 7 | 8 | 9 | 10 |
|---|---|---|---|---|---|---|---|---|---|---|
| A | MeOH blank | 0.25 mg/ml **Cmp. 1** | 0.25 mg/ml **Cmp. 2** | 0.125 mg/ml **Cmp. 1** | 0.125 mg/ml **Cmp. 2** | 0.0625 mg/ml **Cmp. 1** | 0.0625 mg/ml **Cmp. 2** | 0.25 mg/ml **Cmp. 3** | 0.125 mg/ml **Cmp. 3** | 0.0625 mg/ml **Cmp. 3** |
| B | MeOH blank | 1:2 | 1:2 | 1:2 | 1:2 | 1:2 | 1:2 | 1:2 | 1:2 | 1:2 |
| C | MeOH blank | 1:4 | 1:4 | 1:4 | 1:4 | 1:4 | 1:4 | 1:4 | 1:4 | 1:4 |
| D | MeOH blank | 1:8 | 1:8 | 1:8 | 1:8 | 1:8 | 1:8 | 1:8 | 1:8 | 1:8 |
| E | plant medium blank | 1:16 | 1:16 | 1:16 | 1:16 | 1:16 | 1:16 | 1:16 | 1:16 | 1:16 |
| F | plant medium blank | 1:32 | 1:32 | 1:32 | 1:32 | 1:32 | 1:32 | 1:32 | 1:32 | 1:32 |
| G | plant medium blank | 1:64 | 1:64 | 1:64 | 1:64 | 1:64 | 1:64 | 1:64 | 1:64 | 1:64 |
| H | plant medium blank | 1:128 | 1:128 | 1:128 | 1:128 | 1:128 | 1:128 | 1:128 | 1:128 | 1:128 |

*Table S 3. Number of germinated seeds per well in the 96 well plate herbicidal assay*

| | 1 | 2 | 3 | 4 | 5 | 6 | 7 | 8 | 9 | 10 |
|---|---|---|---|---|---|---|---|---|---|---|
| A | >5 | 0 | 0 | 0 | 0 | 0 | 0 | 0 | 1 | 2 |
| B | >5 | 0 | 1 | 0 | 3 | >5 | >5 | 0 | 4 | >5 |
| C | >5 | 0 | 4 | 1 | >5 | >5 | >5 | >5 | >5 | >5 |
| D | >5 | 3 | >5 | >5 | >5 | >5 | >5 | >5 | >5 | >5 |
| E | >5 | >5 | >5 | >5 | >5 | >5 | >5 | >5 | >5 | >5 |
| F | >5 | >5 | >5 | >5 | >5 | >5 | >5 | >5 | >5 | >5 |
| G | >5 | >5 | >5 | >5 | >5 | >5 | >5 | >5 | >5 | >5 |
| H | >5 | >5 | >5 | >5 | >5 | >5 | >5 | >5 | >5 | >5 |

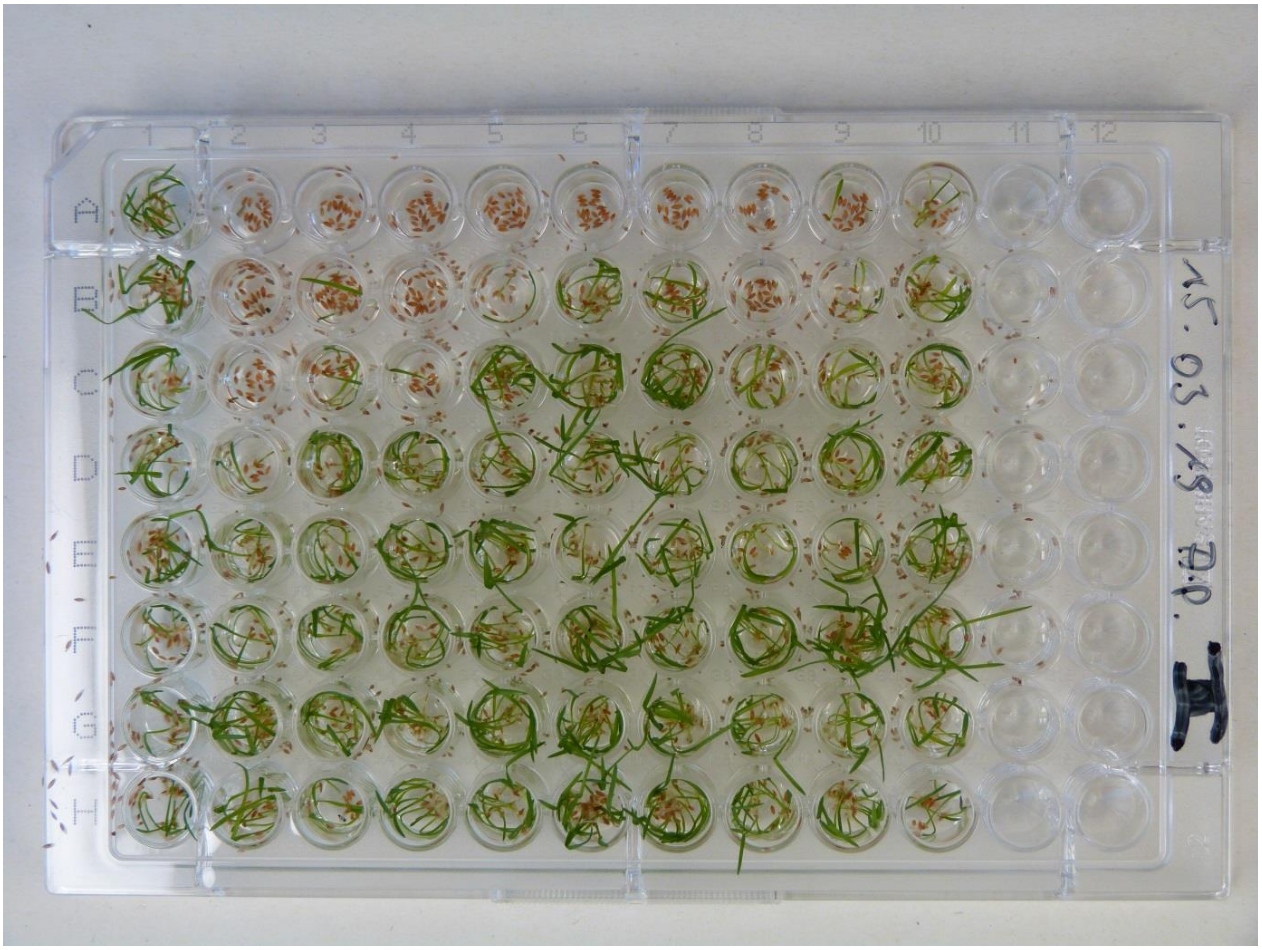

*Figure 3.* Agrostis stolonifera *penncross based Herbicide assay*

### 7 NMR spectra of natural 3,5 dibromo p-anisic acid

This section contains the NMR spectra of the 3,5 dibromo p-anisic acid isolated from the planctomyces strain 10988 in MeOH-*d4.*

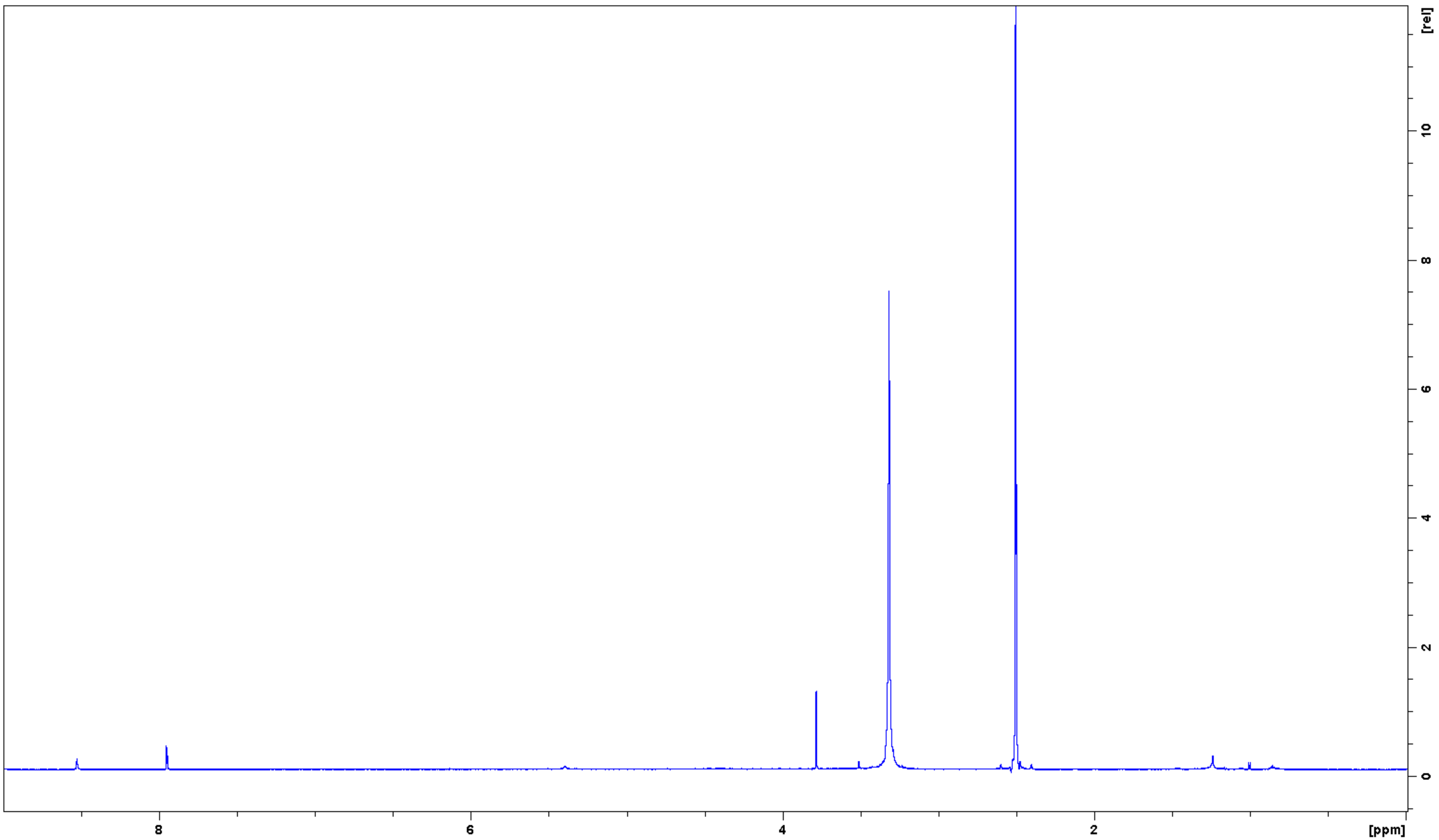

*Figure 4. [1]H NMR spectrum of isolated 3,5 dibromo p-anisic acid in MeOH-d4*

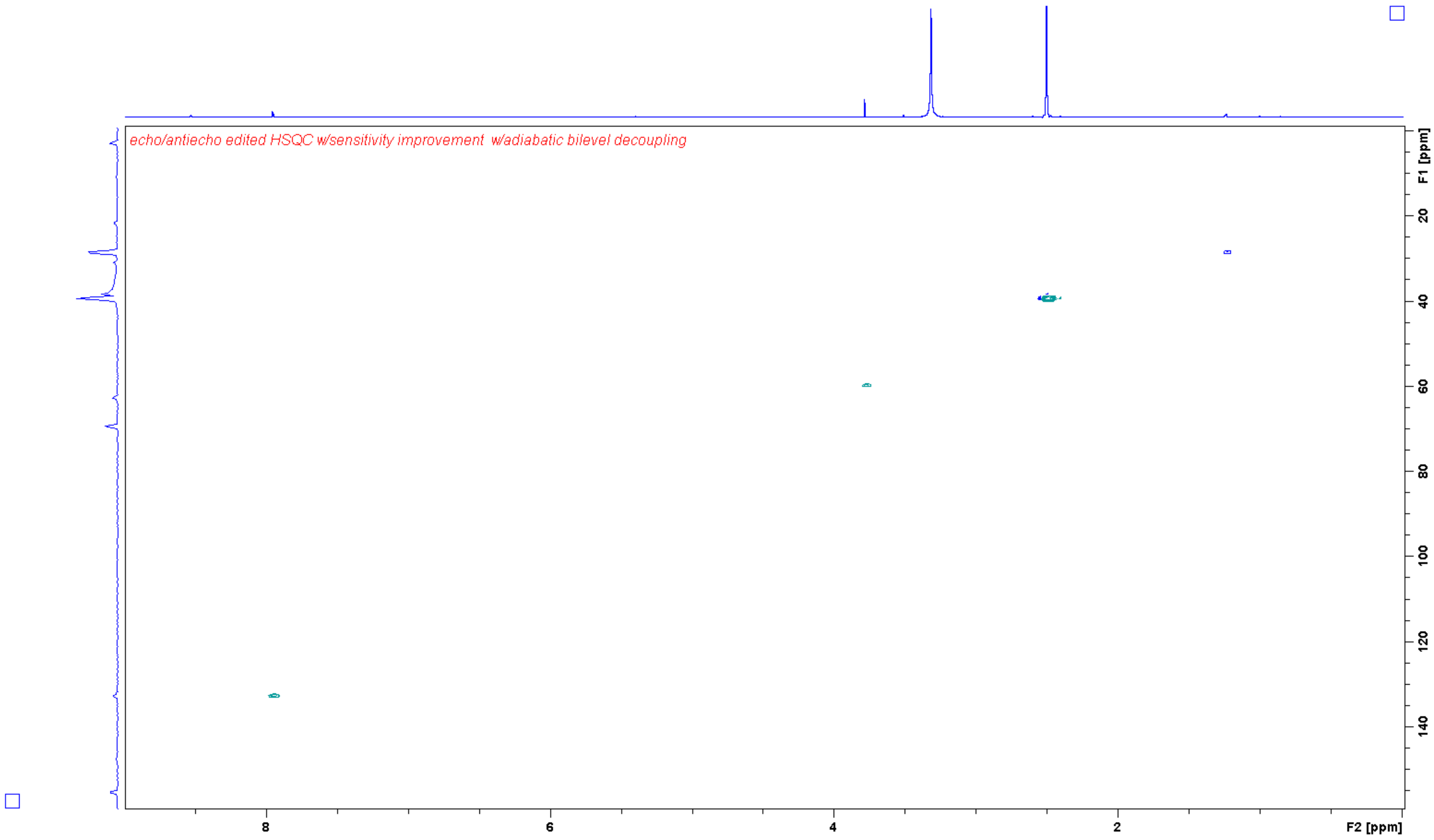

*Figure 5. HSQC spectrum of isolated 3,5 dibromo p-anisic acid in MeOH-d4*

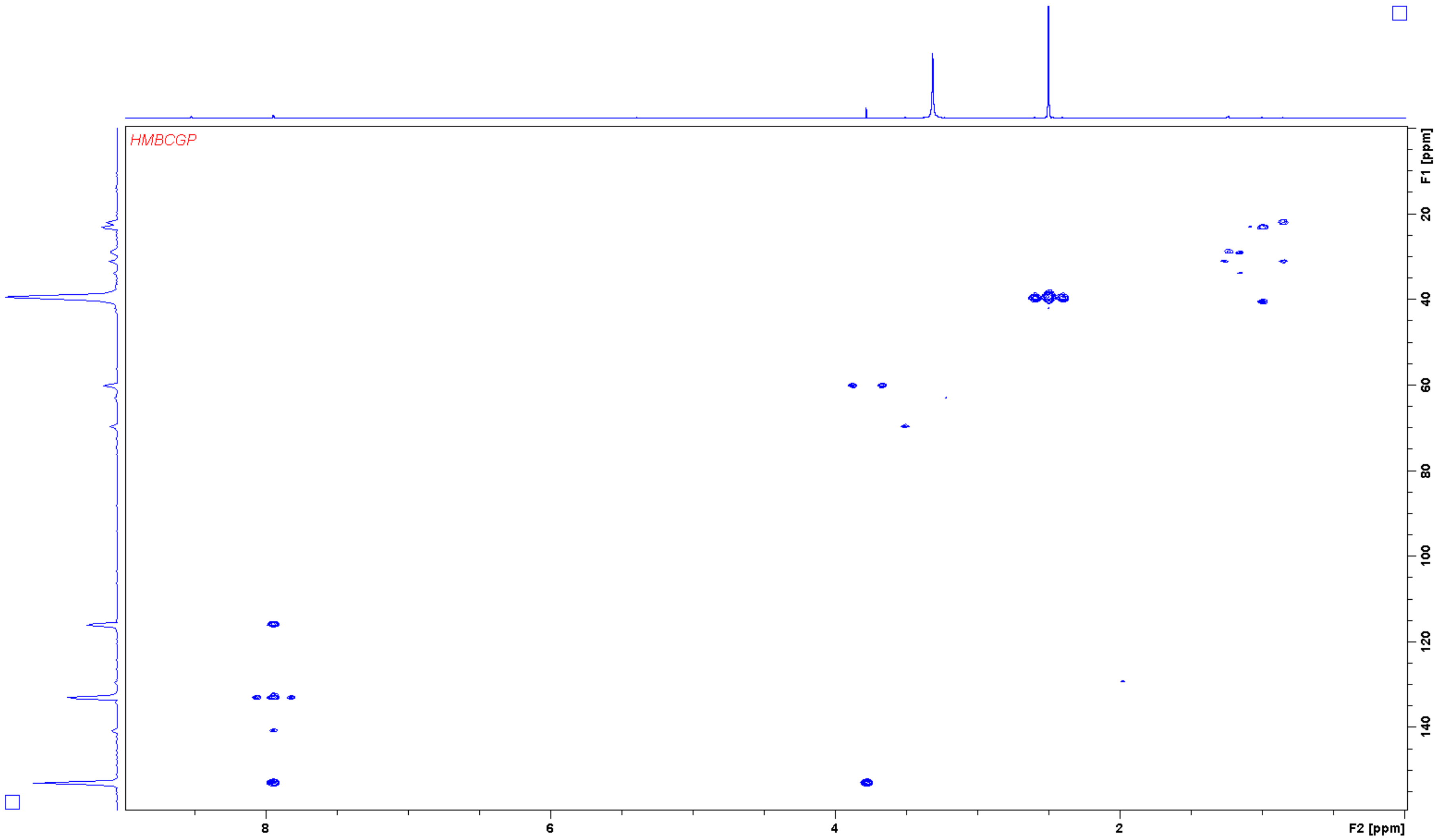

*Figure 6. HMBC spectrum of isolated 3,5 dibromo p-anisic acid in MeOH-d4*
